# Adversarial Deep Structural Networks for Mammographic Mass Segmentation

**DOI:** 10.1101/095786

**Authors:** Wentao Zhu, Xiaohui Xie

## Abstract

Mass segmentation is an important task in mammogram analysis, providing effective morphological features and regions of interest (ROI) for mass detection and classification. Inspired by the success of using deep convolutional features for natural image analysis and conditional random fields (CRF) for structural learning, we propose an end-to-end network for mammographic mass segmentation. The network employs a fully convolutional network (FCN) to model potential function, followed by a CRF to perform structural learning. Because the mass distribution varies greatly with pixel position, the FCN is combined with position priori for the task. Due to the small size of mammogram datasets, we use adversarial training to control over-fitting. Four models with different convolutional kernels are further fused to improve the segmentation results. Experimental results on two public datasets, INbreast and DDSM-BCRP, show that our end-to-end network combined with adversarial training achieves the-state-of-the-art results.

## 1. Introduction

According to the American Cancer Society, breast cancer is the most frequently diagnosed solid cancer and the second leading cause of cancer death among U.S. women. Mammogram screening has been demonstrated to be an effective way for early detection and diagnosis, which can significantly decrease breast cancer mortality [23]. However, screenings are also associated with high false positive rates, high variability among different clinicians, and overdiagnosis of insignificant lesions [23]. To address these issues, it is important to develop fully automated mammographic image analysis tools that can increase detection rate while reducing false positives.

There are two types of mammographic abnormality for most abnormal cases: micro-calcifications and masses [24]. Micro-calcifications range from 20 to several hundred microns in diameter and typically cluster together. On the other hand, masses are featured by different sizes, shapes, margins (the border of masses), and associated findings (i.e. architectural distortions). These features are strongly correlated with mass classification or diagnose (benign versus malignant). Mass analysis is more challenging due to the large variation in sizes and shapes, and the low signal-to-noise ratio [30]. Mass segmentation provides these features, such as shape and size, which plays a fundamental role for mass detection and classification.

Traditional studies on mass segmentation rely heavily on elaborate human designed features, which can be divided into four categories based on how the mass is modeled: region-based methods, contour-based methods, clustering-based methods and model-based methods [24]. Region-based methods use homogeneous or connective constraint in the mass region generation [10]. Contour-based methods typically need complex image processing or mathematical models to detect boundaries using methods such as level sets [2, 4]. Clustering-based methods group pixels with similar features and need additional post-processing step to eliminate the non-connected regions [28]. Model-based methods build classifiers and learn the intrinsic features from the mass [3].

Inspired by the successful use of deep features in natural image processing [33], in this paper, we focus on using the convolutional neural networks (CNN) to learn features for mammographic images. For deep neural networks, there are two main challenges in mammographic image analysis. The first is the scarcity of publicly available mammogram data which can lead to heavy over-fitting on the training set. Several works have combined different models together to reduce the over-fitting [8, 9]. However, these models use the same objective functions of sub-models, which cannot solve the over-fitting problem fundamentally. The second is the poor contrast of the mammographic images which makes the model predict blurry boundaries and yields inaccurate predictions inside the boundaries. Some works use the structural learning to cope with this difficulty [7, 28]. However, the two-stage training used in these works cannot fully explore the power of potential functions.

To this end, we propose an end-to-end adversarial deep structural network to perform mass segmentation (Fig. 1). The proposed network is constructed by taking advantage of many insights from recent successful networks [19, 32, 11], and is designed to robustly learn from the poor contrastive and scarce mammographic images. Specifically, an end-to-end fully convolutional network (FCN) with CRF structural learning is applied to the mass segmentation. Adversarial training is introduced into the network to learn robustly from the scarce mammographic images. To fully explore the features of mass regions, categorical distributions considering the positions are added for predictions. With these components, we validate our adversarial deep structural network on two public mammographic mass segmentation datasets. The proposed network is shown to consistently outperform other algorithms for mass segmentation.

**Figure 1:**
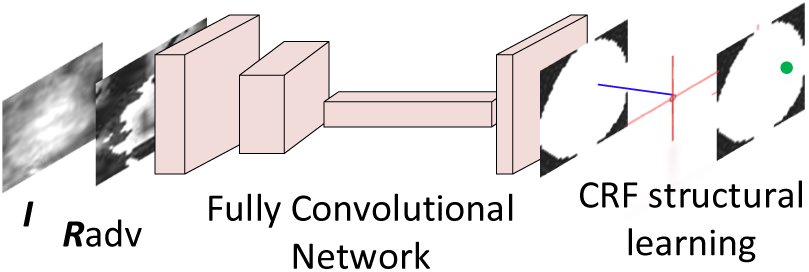
The proposed adversarial deep FCN-CRF network with four convolutional layers followed by CRF structural learning.

Our main contributions in this paper are: (1) Integrating CNN+CRF and adversarial training into a unified end-to-end training framework has not been attempted before. Both components are essential and necessary for achieving the state-of-the-art performance. (2) We employ an end-to-end trained network to do the mass segmentation while previous works needed a lot of hand-designed features or procedures, such as calculating potential functions independently. (3) Our model achieves state-of-the-art results on two commonly used mammographic mss segmentation datasets, the Inbreast dataset and the DDSM-BCRP dataset.

The rest of this paper is organized as follows. Sec. 2 introduces the related work of mammographic mass and natural image segmentation. Sec. 3 describes the framework of adversarial deep FCN-CRF network for mass segmentation. Sec. 4 shows experimental comparisons among these methods. The paper concludes in Sec. 5.

## 2. Related Work

### 2.1. Medical Image Segmentation with Neural Networks

In contrast to the natural image segmentation, only a few works have leveraged neural networks for medical image segmentation. Ngo and Carneiro used deep belief networks as the initialization of distance regularized level set method to segment ventricle in magnetic resonance (MR) images [1]. The deep contour-aware network employed multi-task learning to learn the segmentation and contour of glands hierarchically [5]. Liskowski and Krawiec trained deep convolutional network on a large dataset to do retinal segmentation, and obtained impressive results [18]. The DeepCut network iteratively updated weak annotations and learned the convolutional neural network followed by CRF to do brain and lung segmentation in MR images [26].

Compared to other medical images, mammographic mass segmentation has its specific issues and challenges, such as low signal-to-noise ratio, the poor contrast nature of mass and normal breast tissues. There are few work using deep networks to process the mammogram [7, 34]. Dhungel et al. employed multiple deep belief networks (DBNs), GMM classifier and priori knowledge as the potential functions, and structural SVM (SSVM) to perform the spatially structural learning [8]. They also used CRF with tree reweighted belief propagation to boost the segmentation results [9]. A recent work used the output from a CNN as an additional potential function, yielding the state-of-the-art performance on mass segmentation [7]. However, the two-stage training used in these methods produces potential functions that can easily over-fit the training data, and cannot handle spatial information that might be important for segmentation.

### 2.2. Adversarial Training and Deep Structural networks

Adversarial examples are generated by making small perturbations in the input space that can significantly increase the loss function corresponding to these samples [29]. The adversarial examples can be used to force the model to learn more robust features, which is a novel regularization technique for deep networks and can yield better performance than the base deep networks [11, 21, 20].

Deep structural learning is an emerging topic for natural image processing. The recurrent convolutional network learns unified features of pixels and spatial structures for scene labeling [25]. The fully convolutional network replaces the fully connected layers in CNN with convolutional layers to preserve the spatial structure information [19]. Chen et al. refines the FCN results using the fully connected CRF [15] to do spatially structural learning [6]. The CRF as Recurrent Neural Networks (RNN) further unrolls the fully connected CRF, interprets it as a RNN, and trains the model with FCN followed by CRF as an end-to-end network [32]. Lin et al. further explores CNN as the pairwise potential function in CRF capturing semantic correlations of neighboring patches [17, 16].

The successful models for natural images easily over-fit the training set, which leads to poor performance for mammographic image processing tasks of scarce and low signal-to-noise ratio data. The adversarial training provides effective regularization for deep structure networks when the data is scarce. The end-to-end training scheme eliminates over-fitting of FCN compared with the two-stage training [7]. The priori knowledge varying with pixel positions can be used to enhance the FCN in mass segmentation further.

## 3. Adversarial FCN-CRF network

Leveraging the insights from recent successful deep convolutional networks for natural image segmentation, we design an end-to-end trained fully convolutional network-CRF (FCN-CRF) network with adversarial training. Fig. 1 shows the architecture of the proposed network which has four convolutional layers, and one CRF as RNN layer. For adversarial training, the input to the network is the adversarial examples. To fully consider the characters of mammographic image segmentation, a position priori is integrated into FCN.

In the following section, we first review the FCN-CRF network [32] briefly to make the paper self-contained. We then introduce how to explore the priori knowledge of the mass segmentation and add it into the FCN. Lastly, we describe the adversarial training of the network which enables effective learning for the scarce mammographic images.

### 3.1. Overview of FCN-CRF Network

Fully convolutional network is a successful model applying convolution neural network (CNN) to the image segmentation task [19]. The advantage of FCN over the model with CNN plus fully connected networks is that FCN can preserve the spatial structure of predictions. FCN consists of convolution operator, de-convolution operator [31], or max-pooling in each layer.

For training, the FCN optimizes the maximum likelihood loss function

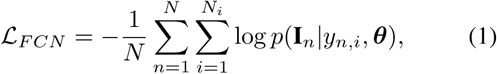

where **I**_*n*_ denotes the *n*th image, *y_n,i_*, is the label of *i*th pixel in the *n*th image, *N* is the number of training mammographic images, *N_i_* is the number of pixels in the image, and ***θ*** is the parameter of FCN. In our case, the size of images is fixed 40 × 40. That is, *N_i_* is 1,600.

CRF is a commonly used method in the structural learning well suited for pixel-wise labeling of image segmentation. In image segmentation, CRF models pixel labels as random variables in a markov random field (MRF) conditioned on a global observation. To make the annotation consistent, we still use *y* = (*y_1_,y_2_,…,y_i_,…,y_16oo_*)^*T*^ to denote the random variables of pixel labels in an image, where *y_i_* ∊ {0,1}. The 0 denotes the pixel belonging to background, and 1 denotes it belonging to mass region. The Gibbs energy of fully connected pairwise CRF [15] is

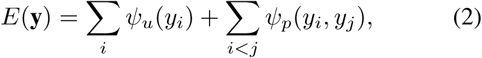

where unary potential function *ψ_u_*(*y_i_*) is the loss of FCN in our case, pairwise potential function *ψ_p_*(*y_i_*, *y_j_*) defines the cost of labeling pair (*y_i_*, *y_j_*).

The pairwise potential function *ψ_p_* (*y_i_*, *y_j_*) can be defined as

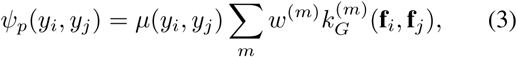

where label compatibility function *μ* is given by the Potts model in our case,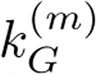 is the Gaussian kernel applied to feature vectors, *w^(m)^* is the learned weight. Pixel values *I_i_* and positions *p_i_* can be used as the feature vector **f**_i_.

Efficient inference algorithm can be obtained by mean field approximation *Q*(*y*) = Π_*i*_ *Q_i_*(*y_i_*) [15]. The update rule is

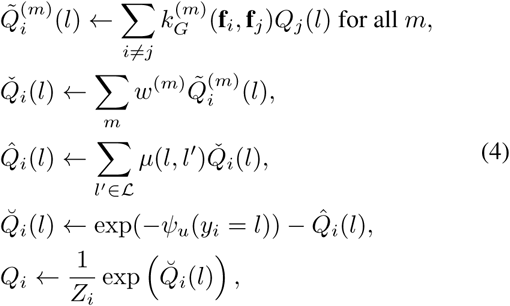

where the first line is the message passing from label of pixel *i* to label of pixel *j*, the second line is re-weighting with the learned weights *w^(m)^*, the third line is compatibility transform, the fourth line is adding unary potentials, and the last step is normalization operator. Here *𝓛* = {0,1} denotes background or mass. The initialization of inference use unary potential function as 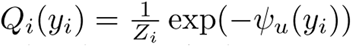

The above mean field approximation can be interpreted as a recurrent neural network (RNN) in Fig. 2 [32]. The recurrent unit takes last state’s approximation *Q_t_*−1, unary potential function *ψ_u_*, and the Gaussian kernel **k***_G_* as the input. The output is updated approximation *Q_t_*. After interpreting the CRF as a RNN, the framework using FCN as potential function followed by CRF can be trained with an end-to-end scheme [32].

**Figure 2:**
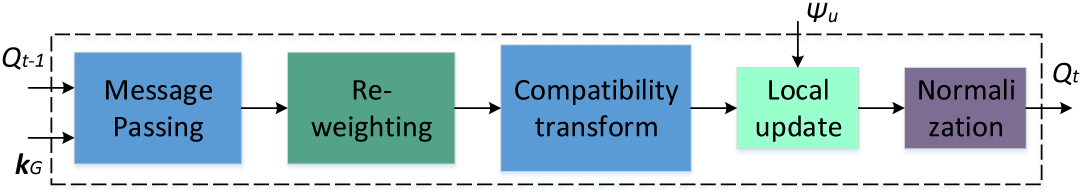
The recurrent unit in the CRF as RNN, which consists of massage passing, re-weighting, compatibility transform, local update and normalization.

### 3.2. FCN Enhancement with Position Priori

The categorical distribution varies greatly with the pixel positions in our mammographic mass segmentation. From observation, most of the mass located in the center of ROI, and the boundary of ROI is more likely background, as shown in Fig. 3. The phenomenon is also validated by the result that priori alone gives much better segmentation than multiple DBNs and GMM classifier [7].

**Figure 3:**
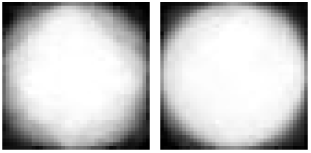
The empirical estimation of priori on INbreast (left) and DDSM-BCRP (right) training datasets.

The conventional FCN provides predictions for pixels inependently. It only considers global class distribution difference corresponding to the number of filters (channels) in the last layer. Here we take the categorical priori of different positions into consideration and add it into the FCN prediction as

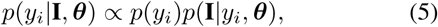

where *p*(*y_i_*) is the categorical priori distribution varied with the pixel position *i*, and *p*(**I**|*y_i_*, *θ*) is the prediction of conventional FCN. The − log*p*(*y_i_* |**I**, *θ*) is used as the unary potential function for *ψ_u_*(*y_i_*) in the CRF as RNN. For multiple FCNs as potential functions, the potential function is defined as

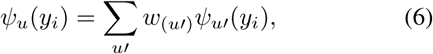

where *w*(_*u′*_) is the learned weight for unary potential function, *ψ_u′_*(*y_i_*) is the potential function provided by one FCN. Experimental result shows that adding position priori obtains 0.25% Dice index improvement than the result of FCN without position priori on the INbreast dataset [22].

### 3.3. Adversarial FCN-CRF Network

Adversarial training provides strong regularization for deep networks [11]. The idea of adversarial training is that if the model is robust enough, it should be invariant to example with the worst perturbation (adversarial examples [29]). The perturbation ***R*** for adversarial example can be obtained as

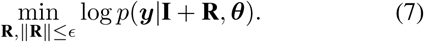

In general, the calculation of exact ***R*** is intractable because the exact minimization is not solvable w.r.t. ***R***, especially for complicated models such as deep networks. The linear approximation and *L*_2_ norm box constrain can be used for the calculation of perturbation [11] as

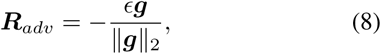

where *g* = ∇_*I*_ log*p*(*y*|***I***, ***θ***). For adversarial fully convolutional network, the network predicts label of each pixel independently as *p*(*y*|***I***, ***θ***) = **Π***_i_ p*(*y_i_*|***I***, ***θ***). For adversarial CRF as RNN, the prediction of network relies on mean field approximation inference as *p*(*y*|***I***, ***θ***) = **Π***_i_ Q*(*y_i_*|***I***, ***θ***).

The adversarial training forces the model to fit examples with the worst perturbation as well. The adversarial loss is defined as

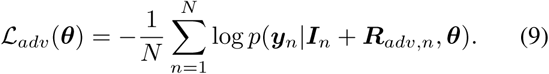

In training, the total loss is defined as the sum of adversarial loss and the empirical loss based on training samples as

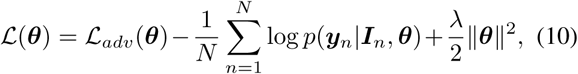

where λ is the regularization factor used to avoid over-fitting, *p*(*y_n_*|***I****_n_*, ***θ***) is either prediction in the enhanced FCN or posteriori approximated by mean field inference in the CRF as RNN for the *n*th image ***I****_n_*. The *L*_2_ regularization term is used only for the parameters in CRF.

### 3.4. Mass Segmentation using the Learned Model

With the learned adversarial FCN-CRF model, the inference *p*(*y_n_*|*I_n_*, ***θ***) is simply the segmentation result of FCN-CRF network because the adversarial training is a regularization of the networks. The inference of FCN-CRF can be calculated directly by running several steps in the CRF as RNN network as Eq. 4. The unary potential functions of CRF as RNN can be obtained as *p(y_i_*|*I*, ***θ***) in Eq. 5. The pairwise potential function of CRF as RNN can be calculated using Eq. 3.

## 4. Experiments

We validate the proposed model on two publicly and most frequently used mammographic mass segmentation datasets: INbreast dataset [22] and DDSM-BCRP dataset [12]. We use the same ROI extraction and re-size principle as [8, 7, 9]. Due to the low contrast of mammographic images, image enhancement technique is used on the extracted ROI images as the first 9 steps of enhancement [2], followed by pixel position dependent normalization is used after the enhancement. The normalization makes the training converge quickly. We further augment each training set by flipping horizontally, flipping vertically, flipping horizontally and vertically, which made the training set 4 times larger than the original training set.

For consistent comparison, the Dice index metric is used for the segmentation results and is defined *Dice* = 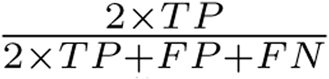. For fair comparison, we also validate the Deep Structure Learning + CNN [7] on our processed data, and obtain similar result (Dice index 0.9013) on the INbreast dataset.

To investigate the impact of each component in our model, we conduct experiments under different configurations defined as follows:

- FCN is the network integrating position priori into FCN. We use the enhanced FCN rather than the conventional FCN in all experiments.
- Adversarial FCN is the FCN with adversarial training.
- Jointly Trained FCN-CRF is the FCN followed by CRF as RNN with an end-to-end training scheme.
- Jointly Trained Adversarial FCN-CRF is the Jointly Trained FCN-CRF with end-to-end adversarial training.
- Multi-FCN is four FCNs with different configurations followed by a linear weighting layer to fuse the four predictions with end-to-end training.
- Adversarial Multi-FCN is the Multi-FCN with end-to-end adversarial training.
- Jointly Trained Multi-FCN-CRF is with the Multi-FCN as potential functions followed by CRF as RNN with end-to-end training.
- Jointly Trained Adversarial Multi-FCN-CRF is the Jointly Trained Multi-FCN-CRF with end-to-end adversarial training.

The configuration of FCN is 6 filters of size 5 × 5 × 1, max pooling of strides 2 × 2 in the first layer, 12 filters of size 5 × 5 × 6, max pooling of strides 2 × 2 in the second layer, 588 filters of size 7 × 7 × 12 in the third layer, 2 filters of size 40 × 40 × 588 for the last deconvolutional layer. Note that we use the configuration in the methods using one FCN. Other three FCNs used in the Multi-FCN are as follows:

- The first layer contains 9 filters of size 4 × 4 × 1, max pooling with strides 2 × 2. The second layer contains 12 filters of size 4 × 4 × 9, max pooling with stride 2 × 2. The third layer has 588 filters of size 7 × 7 × 12. The fourth layer has 2 filters of size 40 × 40 × 588 for deconvolutional operator.
- The first layer contains 16 filters of size 3 × 3 × 1, max pooling with strides 2 × 2. The second layer contains 13 filters of size 3 × 3 × 16, max pooling with strides 2 × 2. The third layer has 415 filters of size 8 × 8 × 13. The fourth layer has 2 filters of size size 40 × 40 × 415 for deconvolutional operator.
- The first layer contains 37 filters of size 2 × 2 × 1, max pooling with strides 2 × 2. The second layer contains 12 filters of size 2 × 2 × 37, max pooling with strides 2 × 2. The third layer has 355 filters of size 9 × 9 × 12. The fourth layer has 2 filters of size size 40 × 40 × 355 for deconvolutional operator.

The parameters of FCNs are set such that the number of each layer’s parameters is almost the same as that of CNN used in the work [7]. For optimization, we use Adam algorithm [13] with learning rate 0.003. The λ used for weights of CRF as RNN is 0.5 in the two datasets. The *∊* used in adversarial training are 0.1 and 0.5 for INbreast and DDSM-BCRP datasets respectively. For mean field approximation or the CRF as RNN, we use 5 iterations/time steps in the training and 10 iterations/time steps in the test phase. Due to the scarcity of data, we simplify the Gaussian kernel *k_G_*(**f**_*j*_, **f**_*j*_) by only considering the neighboring pixels:

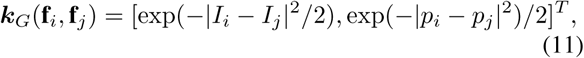

which is a special form of truncated Gaussian kernel.

### 4.1. INbreast Dataset

The INbreast dataset is a recently released mammographic mass analysis dataset, which contains 410 mammographic images [22]. The dataset has 115 cases where 90 cases are from women with both breasts affected. Four types of lesions are included: masses, calcifications, asymmetries and distortions. Compared to other datasets such as mini-MIAS [27], the INbreast dataset provides more accurate contours of lesion region and the mammographic images are of high quality.

For mass segmentation, the dataset contains 116 mass regions. We use the first 58 masses for training and the rest for test, which is of the same protocol used in these works [8, 7, 9].

We have compared our schemes with other recently published mammographic mass segmentation methods [4, 8, 9, 7]. We summarize the results in terms of the publicly used Dice index in Table 1.

**Table 1:** Comparisons on INbreast dataset.

| Method | Dice(%) |
| --- | --- |
| Cardoso et al. [4] | 88 |
| Deep Structure Learning [8] | 88 |
| TRW Deep Structure Learning [9] | 89 |
| Deep Structure Learning + CNN [7] | 90 |
| FCN | 89.48 |
| FCN with Adversarial Training | 89.71 |
| Jointly Trained FCN-CRF | 89.78 |
| FCN-CRF with Adversarial Training | 90.07 |
| Multi-FCN | 90.47 |
| Multi-FCN with Adversarial Training | 90.71 |
| Jointly Trained Multi-FCN-CRF | 90.76 |
| Multi-FCN-CRF with Adversarial Training | **90.97** |

Table 1 shows that the successfully used CNN features in natural image provide superior performance on medical image analysis, outperforming contour based method [4]. Our enhanced FCN achieves about 0.5% improvement than the CNN followed by fully connected network (88.98%). The adversarial training yields 0.3% improvement on average. Incorporating the spatially structural constraint further produces 0.3% improvement. Using model average or multiple potential functions contributes the most to segmentation results which is consistent with work showing that the best model requires five different unary potential functions [7]. Combining all the components together achieves the best performance with relative 9.7% improvement. In our experiment, there is a phenomenon that the FCN over-fits heavily on the training set which can even achieve above 98.60% Dice index. The phenomenon explains why the two-stage training cannot boost the performance too much. The adversarial training works effectively as a regularization to eliminate the over-fitting. We believe the over-fitting is mainly caused by the scarce mammographic images and we strongly support the creation of a large mammographic analysis dataset to accelerate mammographic image analysis research.

### 4.2. DDSM-BCRP Dataset

The DDSM dataset is the largest mammographic image analysis dataset, which contains 2,620 cases (10,480 images) [12]. However, many of images in the DDSM dataset are too old and even damaged. A subset of the DDSM dataset, DDSM-BCRP, is commonly used for its relatively high resolutions and pixel level groundtruth annotations. The DDSM-BCRP dataset includes 89 cases (356 images) for training and 90 cases (360 images) for testing, which contains 39 mass cases (156 images) for training and 40 mass cases (160 images) for testing. After ROI extraction, there are 84 ROI for training, and 87 ROI for test. Comparisons with contour based method [3] and two stage trained deep structure models [8, 9, 7] are summarized in Table 2.

**Table 2:** Comparisons on DDSM-BCRP dataset.

| Method | Dice(%) |
| --- | --- |
| Beller et al. [3] | 70 |
| Deep Structure Learning [8] | 87 |
| TRW Deep Structure Learning [9] | 89 |
| Deep Structure Learning + CNN [7] | 90 |
| FCN | 90.21 |
| FCN with Adversarial Training | 90.78 |
| Jointly Trained FCN-CRF | 90.97 |
| FCN-CRF with Adversarial Training | 91.03 |
| Multi-FCN | 91.17 |
| Jointly Trained Multi-FCN | 91.20 |
| Multi-FCN-CRF jointly training | 91.26 |
| Multi-FCN-CRF with Adversarial Training | **91.30** |

Table 2 shows that, using enhanced FCN achieves comparable performance as the previous best performance. The reason is that there are not many sharp boundaries in the segmentation groundtruth of DDSM-BCRP dataset compared to that of INbreast dataset, which means the variation of groundtruth is much smaller. The position priori contributes more to the model in such case. Adversarial training improves more than 0.5% compared to the model using FCN only. The FCN-CRF with adversarial training obtains about 0.8% improvement over the baseline. The multi-FCN-CRF with end-to-end adversarial training achieves 1.3% (relative 13%) improvement over the Deep Structure Learning + CNN [7]. The result shows the advantage of our proposed network.

### 4.3. Discussions

To further understand the adversarial training, we visualize the segmentation results of FCN (the first row) and FCN with adversarial training (the second row) on the INbreast (a) and DDSM-BCRP (b) datasets as in Fig. 4. For clarity, we only visualize the perimeters of mass prediction results. We randomly select seven samples from the two datasets respectively.

**Figure 4:**
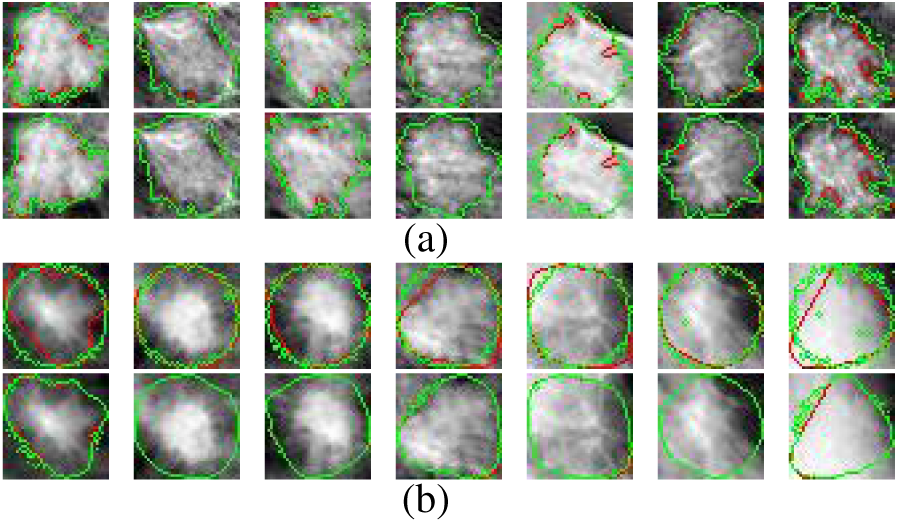
Visualization of segmentation results using the FCN (the first row) and FCN with adversarial training (the second row) on the training sets of INbreast (a) and DDSM-BCRP (b) datasets. Each column denotes different samples. Red lines denote the ground truth. The green lines or points denote the segmentation results.

From Fig. 4, we observe that the FCN with adversarial training yields more accurate borders (green lines) than that of FCN. The FCN with adversarial training obtains better segmentation results than using FCN only on the two datasets. This supports using adversarial training as an effective regularizer to learn better models.

We further visualize segmentation results using the four networks, FCN (the first row), FCN with adversarial training (the second row), jointly trained FCN-CRF (the third row) and FCN-CRF with adversarial training (the fourth row) on the INbreast (a) and DDSM-BCRP (b) datasets in Fig. 5.

**Figure 5:**
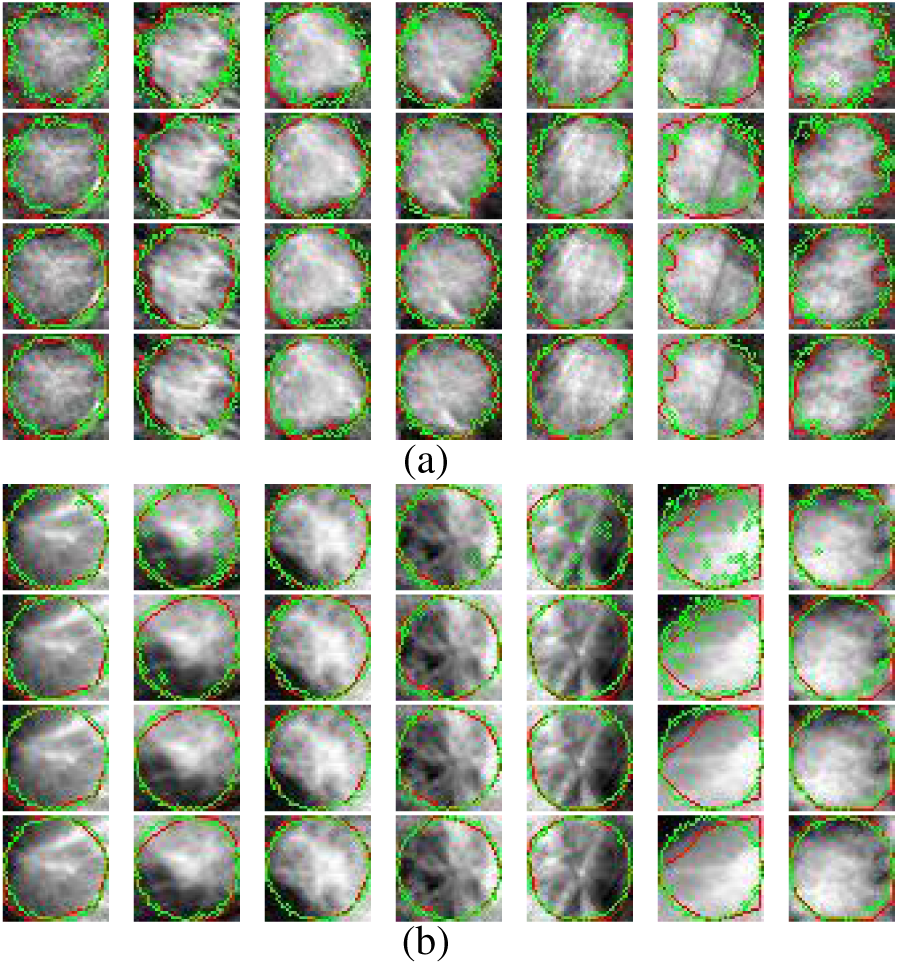
Visualization of segmentation results using the FCN (the first row), the FCN with adversarial training (the second row), jointly trained FCN-CRF (the third row) and FCN-CRF with adversarial training (the fourth row) on the test sets in INbreast (a) and DDSM-BCRP (b) datasets. Each column denotes different test samples. Red lines denote the ground truth. The green lines or points denote the segmentation results.

From Fig. 5, we observe that the segmentations in the first row have vague borders and many outliers within the predicted borders. The segmentations in the second row have fewer vague borders and fewer outliers than the predictions in the first row. The results in the last two rows have sharper and more accurate borders than the first two rows. The phenomenon shows that the CRF based methods achieves better segmentations on the test sets. The structural learning using CRF eliminates outliers within borders effectively, which makes better segmentations and more accurately predicted borders. We will quantize the border prediction in the next paragraph.

We further employ the metric based on the trimap to specifically evaluate segmentation accuracy in boundaries [14]. We calculate the accuracies within trimap surrounding the actual mass boundaries (groundtruth). Trimaps on the DDSM-BCRP is visualized in Fig. 6. We calculate the accuracy in five different trimaps on the two datasets. Five different configurations of the trimaps are set as border size from 1 to 5 with the step 1. Accuracy comparisons among FCN, FCN with Adversarial Training, Jointly Trained FCN-CRF, FCN-CRF with Adversarial Training are shown in Fig. 7

**Figure 6:**
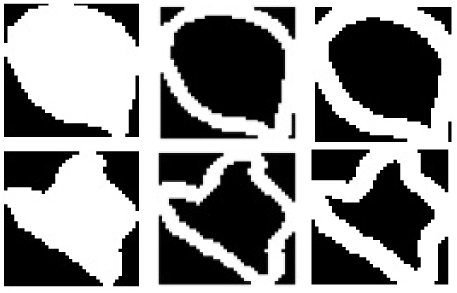
Trimap visualization on the DDSM-BCRP dataset. Images in the first column are the segmentation groundtruth. Images in the second column are the trimaps of width 2. Images in the third column are the trimaps of width 3.

In Fig. 7, the accuracies of FCN-CRF with Adversarial Training are 2-3 % higher than those of FCN-CRF on average and the accuracies of FCN with Adversarial Training are better than those of FCN. The phenomenon show that the adversarial training regularization improves the FCN and FCN-CRF both in the whole image (Dice Index metric) and around the boundaries. The adversarial perturbation is suited to mammographic image segmentation.

**Figure 7:**
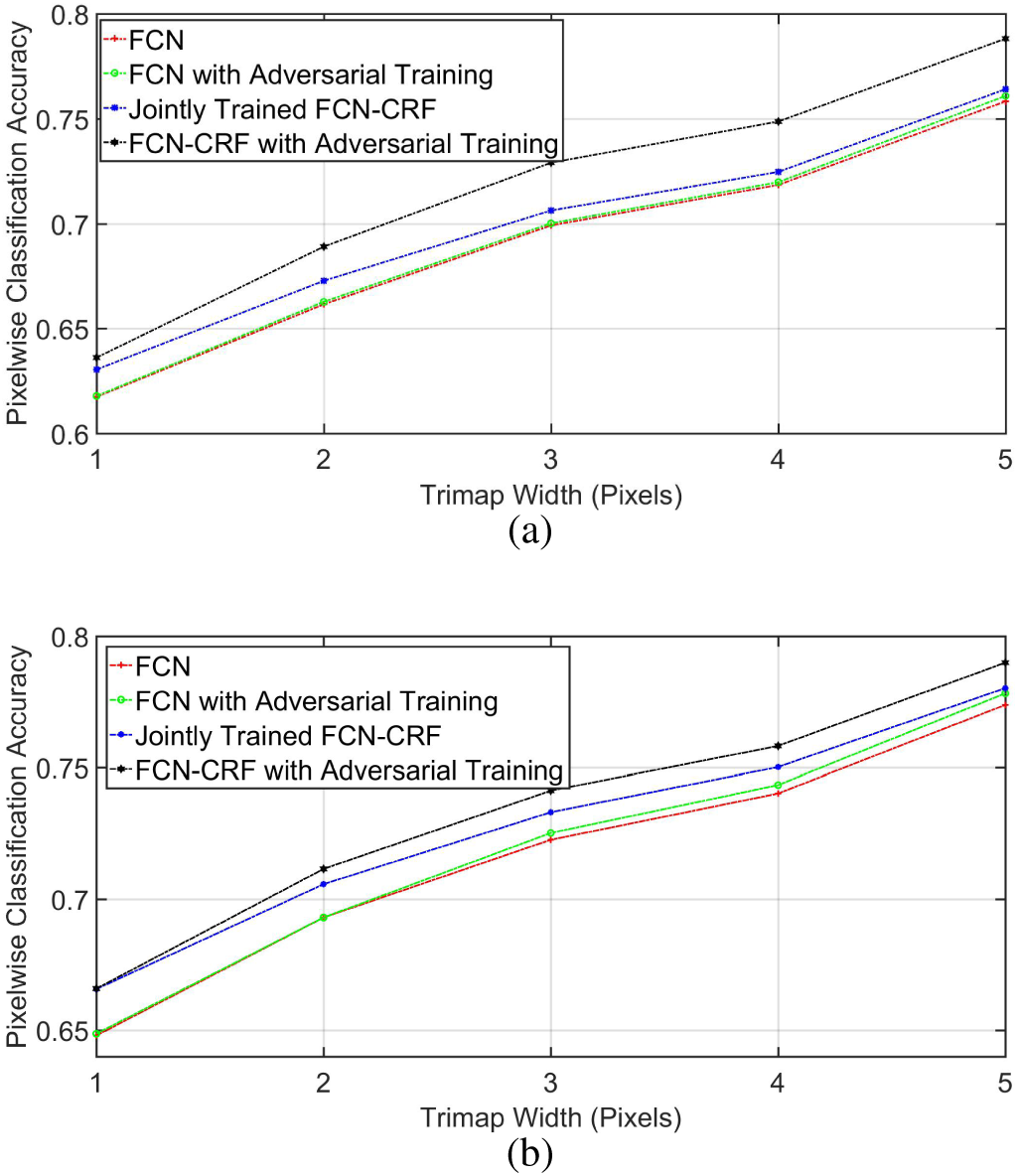
Accuracy comparisons among FCN, FCN with Adversarial Training, Jointly Trained FCN-CRF and FCN-CRF with Adversarial Training in trimaps with pixel width 1, 2, 3, 4, 5 on the Inbreast dataset (a) and the DDSM-BCRP dataset (b). The adversarial training improves the segmentation accuracy on boundaries.

## 5. Conclusion

In this paper, we propose an end-to-end adversarial FCN-CRF network for mammographic mass segmentation. To integrate the priori distribution of masses and fully explore the power of FCN, a position priori is added to the network. Furthermore, adversarial training is used to handle the small size of training data by reducing over-fitting and increasing robustness. Experimental results demonstrate the state-of-the-art performance of our model on the two most used public mammogram datasets.

In future works, it is promising to extend the current work: 1) incorporating multi-scale modeling such as Laplacian pyramid reconstruction to further improve segmentation, and 2) integrating segmentation, mass detection and whole image classification into a unified model with end-to-end training. Our method should be generally applicable to other bio-image analysis problems which are often characterized by the scarcity of training data, frequently small regions-of-interest, and low signal-to-noise ratio.

## Acknowledgments

The authors would like to thank Dr. Neeraj Dhungel from University of Adelaide, Australia for the helpful discussions, the Breast Research Group, INESC Porto, Portugal for the INbreast dataset, and NVIDIA for graphic card donation.

